# Th2 single-cell heterogeneity and clonal interorgan distribution in helminth-infected mice

**DOI:** 10.1101/2021.10.03.462935

**Authors:** Daniel Radtke, Natalie Thuma, Philipp Kirchner, Arif B Ekici, David Voehringer

**Affiliations:** Department of Infection Biology, Universitätsklinikum Erlangen and Friedrich-Alexander University Erlangen-Nürnberg (FAU), 91054 Erlangen, Germany; Institute of Human Genetics, Universitätsklinikum Erlangen and Friedrich-Alexander University Erlangen-Nürnberg (FAU), 91054 Erlangen, Germany

## Abstract

Th2 cells provide effector functions in type 2 immune responses to helminths and allergens. Despite knowledge about molecular mechanisms of Th2 cell differentiation, there is little information on Th2 cell heterogeneity and clonal distribution between organs. To address this, we performed combined single-cell transcriptome and TCR clonotype analysis on murine Th2 cells in mesenteric lymph nodes (MLN) and lung after infection with *Nippostrongylus brasiliensis* (Nb) as a human hookworm infection model. We find organ-specific expression profiles, but also populations with conserved migration or effector/resident memory signatures that unexpectedly cluster with potentially regulatory *Il10*^*pos*^*Foxp3*^*neg*^ cells. A substantial MLN subpopulation with an interferon response signature suggests a role for interferon-signaling in Th2 differentiation or diversification. Further RNA-inferred developmental directions indicate proliferation as a hub for differentiation decisions. We also link long noted *Cxcr3* expression in the Th2 compartment to a population of *Il4*^*pos*^ NKT cells. Although the TCR repertoire is highly heterogeneous, we identified expanded clones and CDR3 motifs. Clonal relatedness between distant organs confirmed effective exchange of Th2 effector cells, although locally expanded clones dominated the response. These results provide new insights in Th2 cell subset diversity and clonal relatedness in distant organs.

## Introduction

Th2 cells are part of the adaptive immune response against helminths and in allergic diseases. They are recruited and differentiate from a pool of naïve CD4 T cells with a wide variety of T cell receptors (TCR) that are formed during T cell development and provide clonotypic specificity to antigens. Differentiated Th2 cells produce the key type 2 cytokines IL-4, IL-5, and IL-13 that elevate type 2 immune responses and thereby promote allergic inflammation but also mediate protection against helminths (Walker & McKenzie, 2018). In recent years, several IL-4 producing Th2 subpopulations have been described and point to substantial heterogeneity within the Th2 population. Only a minor fraction of human IL-4^+^ T cells produces IL-5 which defines them as highly differentiated cells (Upadhyaya, Yin, Hill, Douek, & Prussin, 2011). In the mouse, Th2 cells in the lung generally appear more activated and co-express IL-4 and IL-13 as compared to Th2 cells isolated from lymph nodes of helminth-infected mice (Liang et al., 2011). Th2 cells can further differentiate to follicular T helper 2 (Tfh2) cells that express IL-4, IL-21 and BCL6 and drive humoral type 2 immune responses in the germinal center (GC) (Glatman Zaretsky et al., 2009; King & Mohrs, 2009; Reinhardt, Liang, & Locksley, 2009) although IL-4 secretion by T cells located outside of GCs can be sufficient for GC formation and class switch recombination to IgE (Turqueti-Neves et al., 2014). Tfh13 cells may also develop from Th2 cells in settings or allergic inflammation. These cells co-express IL-4, IL-5 and IL-13 and promote the generation of high affinity anaphylactic IgE in response to allergens (Gowthaman et al., 2019). In addition to these subsets with distinct functions, there are likely different activation and developmental stages present in the Th2 population. Furthermore, fate mapping and adoptive transfer experiments revealed functional plasticity between T helper cell subpopulations which can lead to Th2 cells with remaining or upcoming signatures of other CD4 T cell subsets like Th1, Th9 or Th17 cells (Panzer et al., 2012; Peine et al., 2013; Tortola et al., 2020; Veldhoen et al., 2008).

Infection of mice with the helminth *Nippostrongylus brasiliensis* (Nb) is a widely used model for human hookworm infections with a strong induction of Th2 responses in lung and small intestine (Urban et al., 1992). L3 stage larvae are injected subcutaneously and then first migrate into the lung before they are coughed up, swallowed and reposition to the small intestine where they mature to adult worms. (Urban et al., 1992). Using this model, we have previously shown that Nb infection induces a Th2 response with a broad T cell receptor (TCR) repertoire required for effective worm expulsion (Seidl, Panzer, & Voehringer, 2011). Development of single cell sequencing technology now allowed us to gain a deeper understanding of Th2 cell subsets, TCR clonality and tissue distribution.

Here, we performed single-cell sequencing of T cell receptor (TCR) genes combined with transcriptome profiling of Th2 cells isolated from lung and mesenteric lymph nodes (MLN) at day ten after Nb infection. By this approach, we revealed heterogeneity and differentiation paths within the Th2 compartment, compared Th2 population similarity at distant sites, analyzed cell exchange between organs by clonal relatedness and characterized expanded clones and their TCR sequences.

## Results

### Th2 cells show an organ-specific gene expression profile consistent with acquired effector functions

We performed combined transcriptome and TCR clonotype analysis using the chromium 10xGenomics and Illumina single cell sequencing platform on IL-4-expressingTh2 cells isolated from lung and MLN of two IL-4eGFP reporter (4get) mice (M. Mohrs, Shinkai, Mohrs, & Locksley, 2001) that had been infected 10 days before with Nb (Fig. 1A). IL-4-expressing Th2 cells (CD4^+^IL-4eGFP^+^) were sorted from single cell suspensions of both organs and were directly subjected to scRNA library preparation.

**Figure 1.**
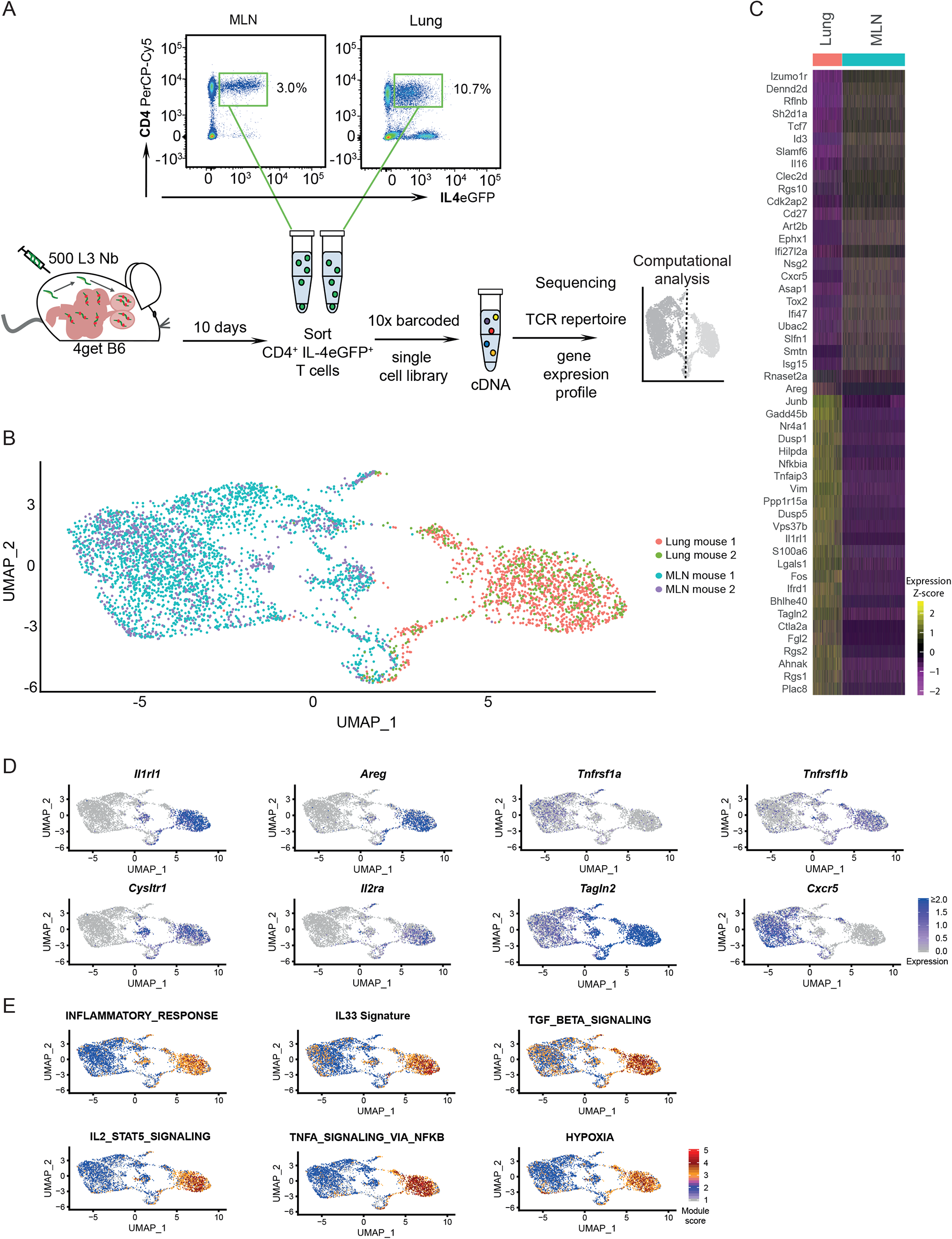
Th2 cells of MLN and lung adopt tissue-specific RNA signatures. A) General experimental outline. MLN and lung cells of two individual Nb-infected IL-4eGFP reporter mice (4get B6) were sorted for IL-4eGFP^+^CD4^+^ cells ten days post infection. Then combined transcriptome and TCR repertoire sequencing was performed. Flow cytrometry plots show the frequency of Th2 cells (IL-4eGFP^+^CD4^+^ cells) in MLN and lung. B) UMAP representation of MLN and lung cells ten days post Nb infection. C) Heatmap of top twenty-five most up- and most down-regulated genes between MLN and lung cells. D) Expression of selected genes or E) gene-signature module scores for single cells on top of UMAP representation.

4get mice were chosen as they allow isolation of Th2 cells *ex vivo* without prior restimulation. In contrast to other IL-4 reporter mice such as the KN2 strain, 4get mice even report the early stages of Th2 differentiation (K. Mohrs, Wakil, Killeen, Locksley, & Mohrs, 2005). Sampling of the lung was performed as Th2 cells accumulate in this organ a few days after Nb infection. Complimentary MLNs were included as a distant secondary lymphoid organ associated to the intestine where the worms reside from day 4 to about day 10 after infection. This setup enabled us to compare Th2 cell subsets and clonotypes in both organs at single cell resolution.

In order to restrict the analysis after sequencing to high quality Th2 cells, we included in total 4710 cells with detected, functional TCR α- and β-chains that also passed our additional QC filters (see methods section) (Suppl. Fig. 1). We used an unbiased high dimensional clustering approach followed by dimensional reduction for simple representation of complex data (Stuart et al., 2019).

Our approach revealed that Th2 cells of lung and MLN have a distinct organ-specific expression profile represented by clear separation of cells from both organs upon dimensional reduction (Fig. 1B) and highlighted by differential expression analysis between lung and MLN cells (Fig. 1C). While most Th2 cells from the MLN (MLN cells) express the gene for the chemokine receptor CXCR5 associated with homing to B cell follicles and recruitment of Tfh cells to germinal centers (Breitfeld et al., 2000; Schaerli et al., 2000), the majority of Th2 cells from the lung (lung cells) and MLN cells that cluster in proximity to lung cells hardly express it (Fig. 1D). Interestingly these cells rather show an increased expression of the gene for TAGLN2 which stabilizes the immunological synapse and is relevant for proper T cell effector function (Na et al., 2015). A stronger effector phenotype of lung cells is also supported by an increase of inflammation signature genes and hypoxia-associated genes in these cells, which are associated with enhanced glycolysis required for late Th2 effector differentiation (Healey et al., 2021; Stark, Tibbitt, & Coquet, 2019) (Fig. 1E).

In line with enhanced effector function, most lung cells and some proximal MLN cells express the gene for the IL-33 receptor ST2 (*Il1rl1*) that recognizes the alarmin IL-33 and induces IL-5 and IL-13 production. Mice that lack IL-33 are not able to effectively clear intestinal helminths, likely due to defects in the T cell and ILC2 compartments (Hung et al., 2013). We also find an elevated gene signature for IL-33-stimulated T cells in the lung, which suggests active signaling via the ST2 receptor. Amongst several stimuli, ST2 can be up-regulated via the IL-2-STAT5 axis (Guo et al., 2009; Meisel et al., 2001) for which we find an elevated expression of the IL-2 receptor (*Il2ra*) and target genes in the lung. In addition, we find an elevated TNF expression signature together with higher expression of the gene *Tnfrsf1b* encoding for Tumor necrosis factor receptor 2 (TNF-R2) that promotes ST2 expression upon TNF binding (Kumar, Tzimas, Griswold, & Young, 1997). IL-33/ST2 mediated signals in turn induce production of amphiregulin in asthma models and we also find the amphiregulin encoding gene *Areg* to be up-regulated in lung cells upon Nb infection. Amphiregulin promotes tissue repair and resolution of inflammation in tissues (Zaiss, Gause, Osborne, & Artis, 2015) but can also re-program eosinophils to develop a fibrosis-driving effector phenotype (Morimoto et al., 2018).

In our Nb infection model, lung cells and a fraction of proximal clustering MLN cells also express genes associated with asthma or involved in pathways targeted by drugs for asthma treatment like *Cysltr1, Plac8* or *Adam8* (Naus et al., 2010; Tibbitt et al., 2019; Trinh, Nguyen, Choi, Park, & Shin, 2019). Lung cells but only few MLN cells in our dataset also show an increased expression of TGFβ target genes, consistent with the described Th2 cell plasticity towards a Th9 phenotype (Veldhoen et al., 2008) which potentially broadens the T effector functions (Fig. 1C-E).

### Conserved expression profiles for migratory and effector/resident memory Th2 cell populations in lung and mesenteric lymph nodes

Next, we combined an analysis of gene expression with unbiased cluster and RNA velocity analysis to define subpopulations and identify potential developmental and differentiation paths. We define two lung, five MLN and a mixed proliferative cluster numbered by size and further described below (Fig. 2A): L1 (basic activated/effector), L2 (migrating), L+MLN (proliferating), MLN1 (basic activated), MLN2 (contains Tfh2), MLN3 (IFN response signature), MLN4 (effector/resident memory like), MLN5 (migrating), MLN6 (innate-like/NKT), MLN7 (myeloid RNA containing Th2).

**Figure 2.**
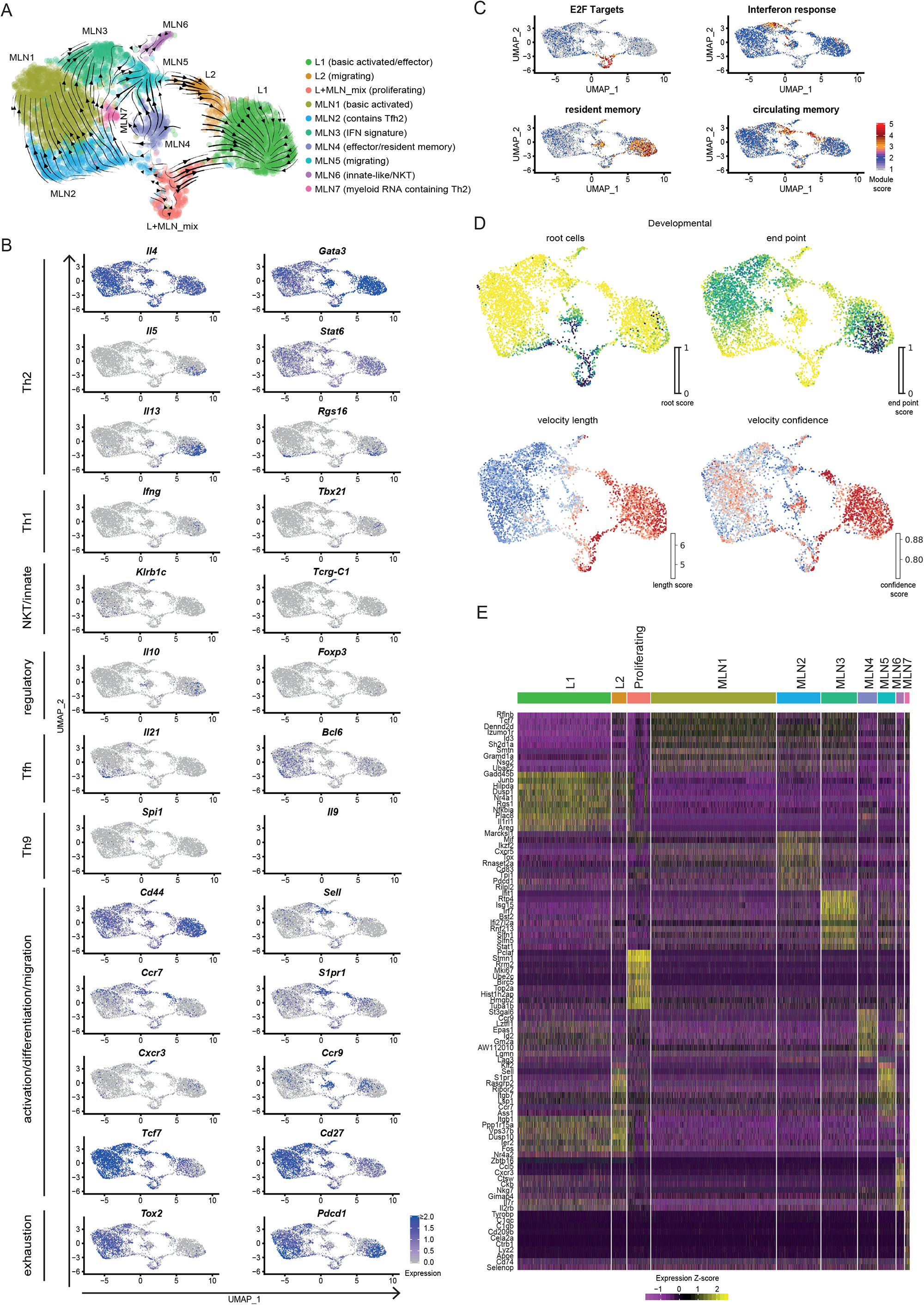
Conserved expression profiles for migratory and effector Th2 populations between organs and their inferred developmental paths. A) *De novo* clustering approach with manually added cell type description. Arrows present developmental paths inferred by RNA-velocity. B) Expression of cell lineage-associated and additional marker genes or C) gene signature scores for single cells on top of UMAPs. D) RNA-velocity defined root cells and developmental end points as well as inferred differentiation speed (velocity length) and velocity confidence for cells are visualized on UMAPs. F) Heatmap of top ten upregulated genes for each cluster compared to all other cells.

We first screened the cells for known markers of T helper cell subsets. The analyzed cells from MLN and lung express the Th2 hallmark genes *Il4, Gata3* and *Stat6* (Fig. 2B). However, only the L1 cluster expresses IL-5 which promotes eosinophil development, recruitment and survival. Similarly, IL-13 is expressed a bit broader in L1 but additionally in MLN4. IL-13 elicits a broad spectrum of effector type 2 immune functions including eosinophilic inflammation, mucus secretion and airway hyperresponsiveness (Rothenberg & Hogan, 2006; Takatsu, Kouro, & Nagai, 2009). According to the pro-inflammatory IL-5 and IL-13 production, double producers are thought to be a strong or pathogenic effector subset of Th2 cells that includes highly differentiated CD27^low^, PD-1(*Pdcd1*)^high^ memory cells (Upadhyaya et al., 2011), which is also reflected on gene expression level in our data. Enhanced *Rgs16* expression of the IL-5^+^/IL-13^+^ cells is also associated with higher cytokine production (Lippert et al., 2003) and further supports effective effector molecule production (Fig. 2B).

As expected very few Th2 cells in L1 express the Th1 hallmark genes *Ifng* and *Tbx21* (encodes T-bet). However, MLN6 expresses the genes for the usually Th1-associated chemokine receptor CXCR3 plus the MLN activation- and lamina propria homing-associated chemokine receptor CCR9 (Campbell & Butcher, 2002; Stenstad et al., 2006). In this population, a fraction of cells also expresses *Zbtb16* which encodes the NKT cell-associated transcription factor PLZF (Savage et al., 2008) and *Klrb1c* (encodes NK1.1) or in addition to the TCRα and TCRβ chains *Tcrg-1c* (encodes the constant region of TCRγ) as an indication that these cells show signs of unsuccessful or not yet successful development into γδ T cells. Hence, MLN6 seems to contain predominantly innate-like / NKT cells (Fig. 2B).

The classical marker for Treg cells FOXP3 was hardly found on gene expression level in our dataset. Nevertheless, small fractions of cells in L1 and MLN4 express *Il10*, which suggests regulatory capacity independent of *Foxp3* expression (Fig. 2B).

T follicular helper cells that express IL-4 (Tfh2) were detected in MLN2 They express both Tfh markers IL-21 and BCL6 on gene level. Of note, *Bcl6* expression seems less restricted to a specific cluster. Where *Il21* and *Bcl6* expression overlaps cells show expression of *Rgs16* associated with enhanced Th2 cytokine production and trafficking (Lippert et al., 2003). In line with a recent publication, we did not observe Tfh13 cells (IL-13^hi^IL-4^hi^IL-5^hi^IL-21^lo^) which were reported to be associated with production of high-affinity anaphylactic IgE in Th2 responses to allergens but not helminth infections (Gowthaman et al., 2019).

Increased expression of TGFβ target genes further suggested Th2 cell plasticity towards Th9 cells in our data set (Fig. 1E). However, we only find very few *Il9* or *Spi1* (encodes the Th9-associated transcription factor PU.1) expressing cells in the lung. *Il9* expression was also barely detectable in the MLN, while *Spi1* was expressed in the MLN4 population (Fig. 2B).

Cluster L2 and MLN5 visually form a “bridge” between the MLN and lung compartments. Indeed cells in these clusters both express genes coding for CD62L (*Sell*), CCR7 and S1PR1 involved in cell adhesion and T cell trafficking suggesting that these clusters contain recent immi-/emmigrants (Fig. 2B). They also express *Tcf7* (associated with self-renewal capacity) and *Cd27* (encoding a central memory T cell marker). In line, the whole “bridge” shows a circulating memory signature (Rahimi, Nepal, Cetinbas, Sadreyev, & Luster, 2020) (Fig. 2C). L2 is the only lung fraction that expresses reasonable amounts of *Cxcr5* and *Tox2, both of which* expressed in a majority of cells in MLN5 (Fig. 1A, 2B). This suggests that the profile of these lung cells in part reflects the profile found in secondary lymphoid organs and strengthens their identification as migrating cells. However, there are also differences between the lung- and MLN-associated “bridge” clusters. Cells in L2 expressed more CD44, which is suggestive of cells in later central memory or effector cell state and also shows more expression of the exhaustion marker encoding *Tox2* and *Pdcd1*. In contrast, MLN5 does not show clear signs of a lung signature (Fig. 2E), potentially reflecting that the visual “bridge” is not a real connection and contains immi-/emmigrants to/from other secondary lymphatic organs like the lung draining lymph nodes or other peripheral organs.

Proliferating cells of both organs have a similar expression profile and fall into the same cluster (L+MLN) as proliferation induces a strong gene signature highlighted by a proliferation-associated E2F signature gene set (Fig. 2C).

As already noted in the comparison between MLN and lung cells on a broad perspective, L1 lung cells seem to have a stronger effector phenotype (e.g. stronger expression of *Il1rl1* (encodes ST2), *Cysltr1* (receptor for cysteinyl-leukotrienes C_4_, D_4_ and E_4_), *Il2ra, Il5, Il13*) but MLN4 has a similar signature. Both populations are high for a published signature of lung resident memory T cells (Rahimi et al., 2020) (Fig. 2C) and for both the effector-associated genes *Plac8* and *Adam8* drive the signature (Fig. 2E and not shown). In contrast to L1, most MLN4 resident memory signature cells express the gene encoding CCR9 relevant for lamina propria homing (Campbell & Butcher, 2002; Stenstad et al., 2006), while lung cells express the gene for CCR8 hardly found in the MLN (not shown). MLN4 and the MLN6 cluster (innate-like/NKT) both express the resident memory signature and *Ccr9*, probably reflecting local effector/resident memory populations that participate in the intestinal immune response. Of note, there is also a *Ccr9*-expressing fraction of cells in L1 with little overlap to the *Il5/Il13* secreting cells (Fig. 2B). These cells might come from or might migrate to the lamina propria. Interestingly, the gene for the exhaustion marker PD-1 is hardly detectable in the MLN but clearly present in the lung. It can be relevant to rescue differentiating T cells from apoptosis under inflammatory conditions (Patsoukis et al., 2015). In contrast, another exhaustion marker-encoding gene, *Tox2* is only marginally expressed in both populations (Fig. 2B).

While Th1 and Th2 cells are often seen as counterparts that can antagonize differentiation of each other, there is also evidence that the Th1 hallmark cytokine IFN-γ promotes proper Th2 heterogeneity and differentiation either directly as suggested by *in vitro* differentiation studies (Wensky, Marcondes, & Lafaille, 2001) or indirectly by inducing activation of DCs with Th2-priming capacity (Connor et al., 2017; Webb et al., 2017). In accordance MLN3 expresses an IFN response signature after Nb infection (Fig. 2C) likely associated with Th2 priming and differentiation. We further identified an unusual subset of Th2 cells (MLN7) which contains genes for the MHCII-associated invariant chain(CD74), complement component C1q, lysozyme and CD209b. Some of these genes are rather associated with myeloid cells like DCs or macrophages. This population might therefore represent cells emulsified with exosomes or RNA containing vesicles during library generation, either externally attached or taken up by the cell.

### RNA-inferred developmental directionality of Th2 cells supports proliferation as a hub for differentiation decisions

To infer developmental relatedness of clusters defined above we performed an RNA velocity analysis in which the ratio of spliced to unspliced RNA transcripts is used to calculate and visualize likely developmental directions (Bergen, Lange, Peidli, Wolf, & Theis, 2020; La Manno et al., 2018) (Fig. 2A). We used the scVelo algorithm (Bergen et al., 2020) which identifies the most undifferentiated cells as root cells and highly differentiated cells as developmental end-points (Fig. 2D), connected by arrows that show likely paths from root to end-points (Fig. 2A). The proliferation cluster (L+MLN) reflects the majority of root cells in our data and highlights proliferation, in accordance with the literature (Gett & Hodgkin, 1998; Gulati et al., 2020; Radtke & Bannard, 2018), as a critical branching point at which differentiation decisions are taken. The MLN part of the “bridge” clusters (MLN5), the IFN signature cluster (MLN3) and the main MLN (Basic activated Th2; MLN1) cluster are marked as relatively diffuse end-points in the MLN, associated with a low differentiation speed and confidence reported by scVelo (Fig. 2D). This suggests that wide parts of the MLN Th2 cells are relatively heterogeneous. The effector/resident memory like cluster (MLN4) is in itself heterogeneous and contains cells with a strong root signature which hardly overlap with the also contained strong resident memory signature cells. A relatively high differentiation speed and confidence compared to other MLN clusters suggests that it contains a fast developing effector/resident memory like population. Based on the MLN5 “bridge” cluster definition as an end-point, it might rather reflect cells that leave the MLN. The lung cluster of the “bridge” (L2) instead contains cells that differentiate with high confidence and inferred speed towards the main lung cluster of effector cells (L1), which suggests that these cells enter the lung and further differentiate locally. The IL-5/IL-13 double producers previously defined as highly differentiated effector cells (Upadhyaya et al., 2011) reflect the end-point in the lung (Fig. 2A,D).

In conclusion, RNA velocity supports proliferation as a hub for differentiation in the Th2 compartment and supports that migratory Th2 cells rather leave secondary lymphatic organs and enter peripheral organs while the reverse migration path is a rare event.

### Clonal relatedness of Th2 cells in distant organs confirms effective exchange of effector cells

The single cell immune profiling approach allows for combined RNA expression profiling and TCR repertoire analysis, which made exploration of clonal relatedness between clusters possible. In line with previous results (Seidl et al., 2011) we find a broad TCR repertoire after Nb infection as the majority of distinct TCRs was only found in one cell. However, 28% of cells expressed a TCR found in at least two different cells (same CDR3 nucleotide sequence, the same variable and joining segments). The most abundant clone has fifteen sequenced members in the samples analyzed which translates to about 8600 estimated members in total lung and MLN tissue. As we only analyze a small sample of the whole organs the calculation clearly underestimates the fraction of expanded T cell clones in the population. A still substantial part of clones is found in both organs, which suggests an effective distribution between MLN and lung. In contrast, only two small clones were identical between the two analyzed mice implicating very few public clones (Fig. 3A). The innate-like/NKT innate cluster (MLN6) and the MLN7 cluster hardly contained expanded clones suggesting limited TCR specificity-driven proliferation in these clusters (Fig. 3B). Cells of the “bridge” clusters (MLN5 and L2) contain substantially more expanded clones but less than the effector/resident memory likepopulations (MLN4 and L1), which in turn contain less expanded clones than the more homogeneous majority of MLN clusters (MLN1, MLN2, MLN3). It might reflect that the “bridge” clusters contain immi-/emmigrated cells from distant sites with less clonal overlap to the local population.

**Figure 3.**
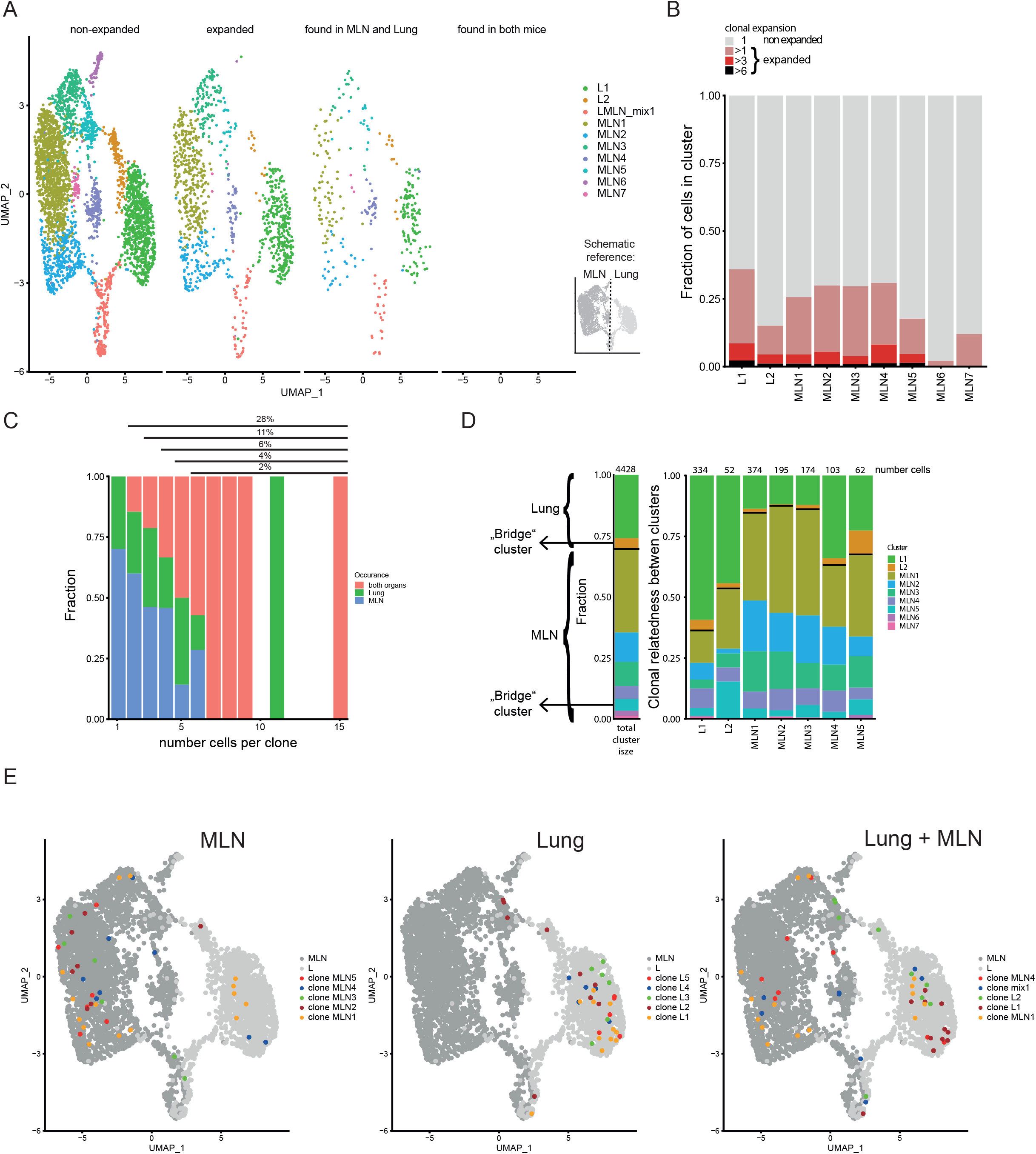
Clonal relatedness of Th2 cells in MLN and lung. A) UMAP of MLN and lung cells split by cells with unique TCRs (non-expanded), cells with the same TCRs found in more than one cell (expanded), cells with the same TCRs found in both organs and cells with the same TCRs found in both mice. Cells are colored by cluster. Schematic drawing roughly highlights how MLN and lung cells are separated on UMAP. B) Stacked bar plot on presence of expanded clones per cluster. Expansion level relates to overall presence in the data set. C) Fraction of cells in MLN, lung and both organs in relation to clone expansion. Numbers above indicate proportion of expanded cells in total population. D) Clonal relatedness between clusters. The stacked bar to the left gives the fraction of each cluster in the data set as a reference (proliferating cluster was excluded). Stacked bar graphs to the right visualize for every expanded clone of a cluster where other members of a clone are found (cluster distribution). Numbers above bars represent the number of cells that each bar represents. Bars for clusters with only few expanded clones are not shown. Black horizontal lines separate the MLN and lung clusters in the bar graphs. Cluster of cells with a migratory signature are highlighted as “bridge cluster”. E) Top five expanded clones by occurrence in MLN (left), lung (center), or in total data set (right).

Next, we find that strongly expanded clones are effectively spread over organs (Fig. 3C). The typical caveats of current single cell technologies (sampling noise and limited sample size) do not allow to draw a similar conclusion for lowly expanded clones (<3 cells per organ). Determination of the clonal relatedness of clusters compared to the overall frequency of a cluster in the data set again highlights effective distribution of effector Th2 cells between distant organs (Fig. 3D). The clones of clusters that are most distant to cells of the other organ (L1, MLN1, MLN2 and MLN3) tend to expand more locally, represented by the higher percentage of related cells found in the same organ compared to the overall distribution of clusters. Directly compared to those clusters, the clones in “bridge” clusters (L2 and MLN5) have a higher frequency of members in the other organ, especially apparent in the other part of the “bridge” in each case. The effector/resident memory like cluster of the MLN (MLN4) also shows increased relatedness to the lung that contains a large number of effector cells in cluster L1. The finding that clusters visualized near the other organ also show enhanced TCR repertoire relatedness to that organ confirms significant inter-organ migration and that the UMAP efficiently displays real relatedness of clusters.

As a next step, we visualized the five most strongly expanded clones determined for each organ separately or we combined counts to get the most strongly expanded clones in the total data set (Fig. 3E). Members of such clones in the MLN tend to be preferentially found in the MLN1 cluster and to a lower extend in the neighboring Tfh-associated cluster (MLN2) and the IFN signature cluster (MLN3). They are hardly found in the lung proximal clusters (MLN4-MLN7). However, few members of four of the top expanded MLN clones are found in the lung and are therefore successfully spread across organs. Top expanded lung clones do not overlap with the top expanded MLN clones and preferentially show up in the big lung cluster in which effector cells are found (L1). In contrast to the MLN only one of the top expanded lung clones has members in the MLN, indicating that these clones successfully expanded locally but have limited capacity to spread to the MLN. The five most highly expanded clones in the whole data set strongly overlap with the ones determined for the separate organs. This indicates that despite remarkable exchange between the distant organs, strong local expanders dominate the response and while more evenly distributed clones are present, they do not outnumber locally expanded ones in a combined analysis of MLN and lung cells.

In summary, there is substantial overlap of expanded clones between the MLN and lung during Nb infection, but rather locally restricted clones successfully expanded in an otherwise diverse pool of Th2 cells.

### No general preference for specific TCR chain compositions

After analysis of single clones in the last part, where we found expansion but no obvious dominant clones, we determined if there are preferentially used TCR segments or segment combinations. First, we included only one representative member per clone and compared if the same combination of TCRα and TCRβ chain segments is shared between the top fifty most frequently used segment combinations in both organs, both analyzed mice and in non-expanded versus expanded clones (Fig. 4A). For the non-expanded clones there was hardly any overlap seen between mice or organs, only two of the fifty combinations were found in three of the four analyzed organs (MLN and lung of two mice). For the expanded clones, there was limited overlap in combined segment usage between organs of one mouse but not the other. Combinations of TCRα- or TCRβ-variable with joining segments and TCRα with TCRβ variable segments also revealed limited overlap in the top used combinations. *Trbv1* was a recurrently used TCRβ variable segment present in frequently used combinations (Suppl. Fig. 2A). Similarly, for single segments there was no obvious preferential usage in expanded clones compared to non-expanded ones. Again, *Trbv1* was one of few constituents that was moderately increased in expanded versus non-expanded clones (Suppl. Fig. 2B-E). In addition, there was no striking difference observed in total CDR3 amino acid length/length distribution that could be indicative for changes in specificity (Davis et al., 1998; Rock, Sibbald, Davis, & Chien, 1994) between expanded and non-expanded clones (Suppl. Fig. 3A). In a finer grained analysis of single TCRα and TCRβ family members, there was also no change in CDR3 length or length distribution (Suppl. Fig. 3B,C). The general TCRα or TCRβ CDR3 length in MLN and lung of the Nb-infected mice is also not altered compared to naive T cells of the peripheral blood (Fig. 4B).

**Figure 4.**
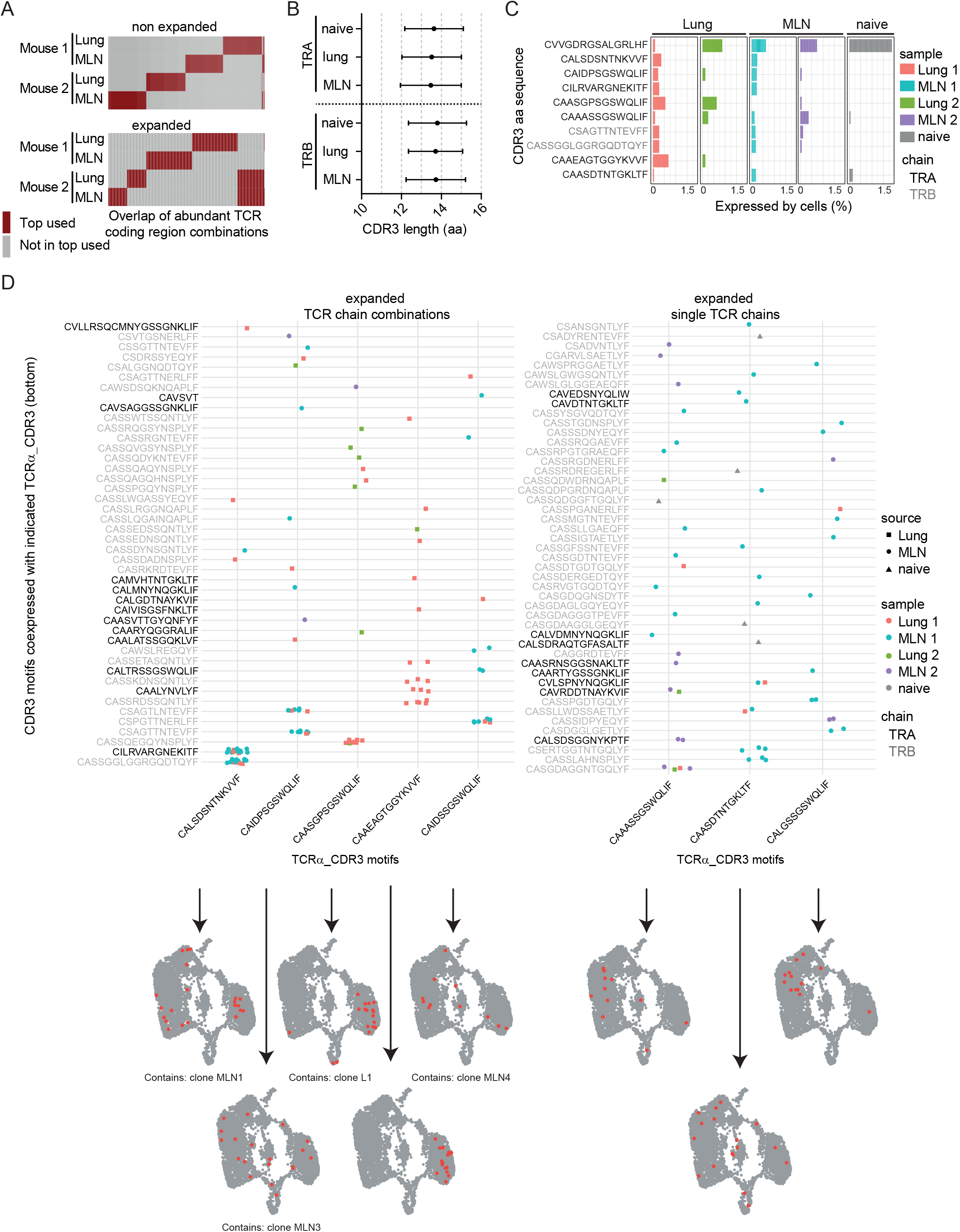
Expanded CDR3 motifs in Th2 cells of Nb-infected mice. TCR repertoire analysis of MLN and lung Th2 cells at day ten post Nb infection. A) Clonotype analysis for overlap of the hundred most commonly used TCR segment combinations (V, J and C region for TCRα; V, D, J and C region for TCRβ) between clonotypes of different mice and organs. Analysis is performed separately for non-expanded and expanded clones. B) Amino acid sequence length of TCRα and TCRβ CDR3 regions. We compare CDR3 regions from peripheral blood T cells of naïve wild-type C57BL/6 mice (naïve) with CDR3 regions of Th2 cells from MLN and lung of Nb-infected mice. C) Most abundant CDR3 amino acid sequences in cells of data set presented as percent of each sample. D) Co-expression of TCRα-related CDR3 motifs (x-axis) with indicated CDR3 motifs of TCRβ or a second TCRα chain in highly expanded clones (left), or highly expanded TCRα_CDR3 sequences in combination with various TCRβ or TCRα chains (right). At the bottom, cells that express the corresponding CDR3 sequences on the x-axis are highlighted on top of UMAP representation of the data set. We also indicate if a CDR3 sequence is associated with the top expanded clones (Fig. 3 E).

In conclusion, expanded clones in the Th2 effector population show no evidence for preferential usage of particular TCRα or TCRβ chains or chain combinations.

### Definition of abundant CDR3 motifs in Th2 cell of Nb-infected mice

T cell antigen-specificity and affinity is mainly confined by variable regions of the TCR and specific T cells are selected and expanded (Rock et al., 1994). We chose the ten most abundant CDR3 sequences on amino acid level that potentially include motifs relevant for anti-helminth immunity and highlight their abundance in different organs and mice (Fig. 4C). TCR data of naïve peripheral blood T cells served as a reference to identify germline-associated CDR3 regions. The most abundant sequence CVVGDRGSALGRLHF was found in all samples (Fig. 4C), including the peripheral blood and represents the CDR3 motif found in the invariant TCRα chain (Vα14-Jα18) of NKT cells (Lantz & Bendelac, 1994). This chain is co-expressed with a variety of different TCRβ chains (not shown) and cells that express such TCRs are primarily found in the innate-like/NKT cluster (MLN6) (Fig. 2A, C). As this cluster also contains some cells with expression of TCRγ chain segments in addition to functional TCRα and TCRβ chains we were able to compare if expression of the invariant TCRα is correlated to TCRγ expression. Indeed, on the one hand, 38% of cells that expressed any TCRγ constant or variable chain segment also expressed the invariant TCRα chain and on the other hand, 62% of cells with the invariant TCRα chain expressed any TCRγ constant or variable chain segment (detection of TCRγ and the invariant TCRα chain in the same cell: correlation 0.48; p<10^−6^). This might suggest a close relatedness of IL-4 expressing γδ T cells with IL-4 expressing NKT cells in a way that cells unsuccessful or not yet successful to generate a functional γδ TCR preferentially develop into αβ NKT cells. Alternatively, NKT cells could induce low level of TCRγ gene expression for other, unknown reasons.

Of the remaining 9 most abundant CDR3 motifs of TCRα or TCRβ chains, seven are not found in the naïve peripheral blood sample (Fig. 4C), which implies an increased probability for them to represent specificity for Nb antigens. Furthermore, only one of these motifs (CAIDPSGSWQLIF) is expressed in both analyzed organs and both mice, which implies that it could be a preferentially selected motif during Nb infection.

We next determined CDR3 motifs that are part of abundant motif combinations (Fig. 4D left panel). As expected, these overlap with the most expanded clones (Fig. 3E) as cells of an expanded clone always use the same chain combination. Only one of the five TCRα CDR3 motifs (CAAEAGTGGYKVVF) was associated with expansion in more than one clone (two clones with same TCRα CDR3 motif but different TCRβ CDR3 motifs). In addition, all five depicted TCRα CDR3 motifs present in abundant pairings are also present in unique pairings with other TCRβ CDR3 motifs. This implies that these motifs are not restricted to an exact TCRα/TCRβ combination or a single clone to be recruited to the Th2 compartment.

As others described (He et al., 2002; Padovan et al., 1993; Padovan et al., 1995) we find T cell clones with expression of two TCRα/TCRβ chain-encoding genes. At least in highly abundant combinations it is unlikely that these are technical artifacts due to contamination with RNA from another cell during library preparation. The clone with the most frequently found combination of CDR3 motifs (clone MLN1) expresses one TCRβ and two TCRα chains, both on average with similar umi counts. Whether both TCRα chains are successfully translated is not known. Of the five depicted TCRα CDR3 motifs, often present in successful CDR3 combinations, four were found in some cells that expressed more than two TCR chains but this frequency is in the expected range for T cells (Alam & Gascoigne, 1998; Balomenos et al., 1995; Davodeau et al., 1995; Dupic, Marcou, Walczak, & Mora, 2019).

In addition to CDR3 motifs found frequently in combinations (expanded clones), we also find abundant CDR3 motifs combined with various other unique TCR chains. (present in several non-expanded clones) (Fig. 4D right panel). These could include CDR3 motifs that provide anti-Nb specificity but failed to induce substantial expansion or accumulation of Th2 cells expressing such TCRs.

In line with a slightly preferential usage of the *Trbv1* gene in expanded compared to non-expanded Th2 cells we find that three of the expanded TCRα CDR3 motifs (CAIDPSGSWQLIF, CAIDSSGSWQLIF, CAASDTNTGKLTF) are preferentially co-expressed with *Trbv1*, which suggest that *Trbv1* is relevant for the immune response against Nb.

## Discussion

Th2 heterogeneity, organ crosstalk and tissue-specific immunity are increasingly appreciated (Schoettler, Hrusch, Blaine, Sperling, & Ober, 2019; Szabo, Levitin, et al., 2019; Szabo, Miron, & Farber, 2019). Here, we applied combined transcriptome and TCR clonotype analysis on Th2 cells across organs upon Nb infection. We identified lung- and MLN-specific gene signatures as well as subpopulations with shared migration and effector/resident memory profiles between organs. We find that expression of tissue damage-associated cytokine coding genes *Il13* and *Il5* is restricted to the effector/resident memory populations in lung and MLN. Interestingly these clusters also contain transcriptionally similar cells that express *Il10* but widely lack expression of the Treg marker encoding gene *Foxp3*. Similar cells have been described in the skin at the infection site after repeated S*chistosoma mansoni* cercaria infection where these cells have immunosuppressive functions (Sanin, Prendergast, Bourke, & Mountford, 2015). Furthermore, effector/resident memory like cells in the MLN are not homogeneous and are found in two clusters of which one is an innate-like/NKT-cluster. Interestingly, the NKT population in this cluster not only co-expresses the invariant NKT cell-associated invariant TCRα chain (Vα14-Jα18) together with a highly diverse repertoire of TCRβ chains but also transcripts for TCRγ chains, which implies shared developmental pathways of NKT and γδ T cells that both tend to express restricted receptor repertoires. The cluster also contains cells that express *Cxcr3*, encoding a typical Th1 marker. CXCR3 has been noted in a small fraction of Th2 cells (Kim et al., 2001) but was not associated with IL-4 producing NKT or γδ T cells before. These findings reveal a heterogeneous effector/resident memory pool in the Th2 population.

When we searched for activation signatures, we found a population of cells with an IFN response signature present in the MLN. Murine *in vitro* studies imply that IFN signaling is needed for proper Th2 differentiation (Wensky et al., 2001). Therefore, Th2 cells with an IFN response signature probably reflect cells that undergo priming or differentiation. Our unbiased RNA velocity analysis further defines them as a differentiation endpoint, suggesting that the IFN response signature is rather related to terminal differentiation. However, the velocity algorithm tries to start from the most undifferentiated cells, which are not necessarily the recently activated and recruited early Th2 cells. Therefore, RNA velocity data needs thoughtful interpretation. However, a fraction of proliferating cells shows low expression of Th2 signature genes and is likely in a stage where differentiation can be determined. In addition to potential developmental paths, RNA velocity analysis suggests that development is faster and has a stricter directionality in the lung compared to the MLN, consistent with the view that the majority of Th2 cells in MLN are less terminally differentiated.

Cells in the migratory clusters of both organs show a weaker organ-specific separation after dimensional reduction but rather form a “bridge” on the UMAP that suggests effective exchange between organs. In line, TCR analysis of our Th2 cells revealed effective exchange of expanded clones between organs. However, the most expanded clones in the one organ were not the most expanded in the other organ. This might relate to different immunological preferences in different compartments or to the different larval stages in which Nb is present in lung and MLN,. Of note, comparison of human bulk TCR repertoires between the lung and its draining lymph node also showed a higher intra-organ TCR repertoire overlap than between organs. This was interpreted to mean that the T cells originate from different precursor pools and recognize distinct antigens (Schoettler et al., 2019). Our data clearly refines that there is effective spread of Th2 effector T cells from the same pool of cells even across distant organs.

Another factor that likely protects from an overshooting Th2 response and could be subject to modulation by the worm is the expression of TNFR2 as impaired TNFR2 signaling leads to augmented Th2 responses (Li et al., 2017). We observed transcripts for TNFR2 preferentially in the lung compared to the MLN after Nb infection, which has not yet been described to our knowledge.

In summary, combined transcriptome and TCR clonotype analysis at single cell resolution provides information about Th2 heterogeneity across organs and reveals relatedness of IL-4 producing NKT cells to γδ T cells. RNA-velocity combined with knowledge from published data appears compatible with a model, in which poorly differentiated proliferating Th2 cells are at a decision-point in their development, with IFN signaling being involved in diversification and differentiation of the Th2 compartment. Despite efficient exchange of expanded Th2 clones between distant organs, the most abundant clones seem to expand locally. Further functional characterization of expanded TCR clonotypes by generating TCR-transgenic mice will help to investigate Th2 cell differentiation, plasticity and memory formation in response to a natural helminth pathogen *in vivo*.

## Materials and Methods

### Mice and *Nippostrongylus brasiliensis* Infection

Two IL-4eGFP reporter (4get) mice (Mohrs et al. 2001) at the age of 13 weeks were infected with Nb. For this, third-stage larvae (L3) were washed in sterile 0.9% saline (37°C) and 500 organisms were injected subcutaneously (s.c.) into mice ten days before analysis. To avoid bacterial infections mice received antibiotics-containing drinking water (2 g/l neomycin sulfate, 100 mg/l polymyxin B sulfate; Sigma-Aldrich, St. Louis, MO) for the first 5 days after infection. Mice were kept under specific pathogen free conditions and were maintained in the Franz-Penzoldt Center in Erlangen. All experiments were performed in accordance with German animal protection law and European Union guidelines 86/809 and were approved by the Federal Government of Lower Franconia.

### Single cell RNA and TCR sequencing

At day 10 after Nb infection lungs and MLNs of IL-4eGFP reporter (4get) mice were harvested. Lungs were perfused with PBS, cut into small pieces and digested with the commercial “Lung Dissociation kit” (Miltenyi, Bergisch Gladbach, GER) according to manufacturer’s instructions. Digested lungs and complete MLNs were gently mashed through a 100 μm cell strainer. For lung cells a 40% percoll purification was applied and erythrocytes were lysed with ACK-buffer (0.15 M NH4Cl, 1 mM KHO3, 0.1 mM Na2EDTA). Then samples of both organs were treated with Fc-receptor blocking antibody (anti-CD16/32, clone 2.4G2, BioXCell, West Lebanon, NH) and stained with anti-CD4-Percp-Cy5.5 antibody (clone: RM4-5). IL-4eGFP^+^CD4^+^ cells were sorted and for each sample, 5000 cells were subjected to 10x Chromium Single Cell 5′ Solution v2 library preparation using the TCR-specific VDJ library kit according to manufacturer’s instructions (10xGenomics, Pleasanton, CA). Gene expression libraries were sequenced on an Illumina HiSeq 2500 sequencer using the recommended read lengths for 10x Chromium 5’ v2 chemistry to a depth of at least 30000 reads per cell. VDJ libraries were sequenced as paired 150 bp reads to a depth of at least 30000 reads per cell.

### Computational analysis

We used Cell Ranger (10x Genomics) to demultiplex sequencing reads, convert them to FASTQ format with mkfastq (Cell Ranger 2.1.1), align them to the murine genome (mm10 v3.0.0) and obtain TCR VDJ clonotypes, consensus sequences, contigs and CDR3 regions (Cell Ranger 3.0.1). TCR associated genes (VDJ and constant region genes for α, β, γ, δ chains) were excluded but kept as metadata to avoid clustering by TCR genes. To be included, cells needed to be defined as such by Cell Ranger and to have > 500 UMIs, > 500 genes detected per cell, < 7% mitochondrial reads and a novelty >0.8 (log10 of gene number divided by log10 of UMIs). Data normalization, differential expression analysis, clustering (based on top 2000 highly variable genes) and dimensional reduction (UMAP based on top 15 principal components) were performed in Seurat (version 3.1.1) (Stuart et al., 2019) under R (version 3.5.1). Gene set-scores for each cell were calculated in Seurat as published before (Tirosh et al., 2016). Gene sets were taken or generated from the published data: resident memory and circulating memory (Rahimi et al., 2020), IL-33 signature (Morimoto et al., 2018), other sets were from the “Molecular Signatures Database” (Subramanian et al., 2005). TCR info was added as metadata to Seurat for combined clonotype and RNA-profile analysis. For RNA velocity, sequencing reads were aligned with kallisto/bustools (version 0.46.2/0.40.0) (Bray, Pimentel, Melsted, & Pachter, 2016; Melsted et al., 2021) to a genome reference with unspliced and spliced RNA variants included (version GRCm39). Obtained information was used as input for scVelo (version 0.2.3) (Bergen et al., 2020) under python (version 3.8.5). UMAP information from Seurat was transferred to scVelo for consistency. Usage of TCR chains and TCR chain-combinations was calculated under R with custom scripts. For TCR/CDR3 analysis, we used the Cell Ranger output and followed a recently developed workflow (according to the “CellaRepertorium” R package) with minor modifications. Contigs that missed a sanity-check were excluded (needed to be productive, full length, high confidence, supported by >1 UMI, CDR3 length > 5 amino acids). Similar CDR3 sequences were not combined (not assuming similar specificity for similar sequences) to maintain higher accuracy. We kept all TCR chains of T cell clones with two TCRα/TCRβ chain-encoding genes expressed for the same reason. Data is available via GEO (GSE181342) and the 10xGenomics TCR reference data set via the 10xGenomics website: PBMCs from C57BL/6 mice (v1, 150 x150), Single Cell Immune Profiling Dataset by Cell Ranger 3.0.0, 10x Genomics, (2018, November 19).

## Acknowledgments

We thank Kilian Schober for helpful discussion. This work was supported by the Deutsche Forschungsgemeinschaft (DFG) with grants RTG1660, FOR2599_TP4, and CRC1181_A02 to D.V.

## Supplementary Information for

**Suppl. Figure 1.**
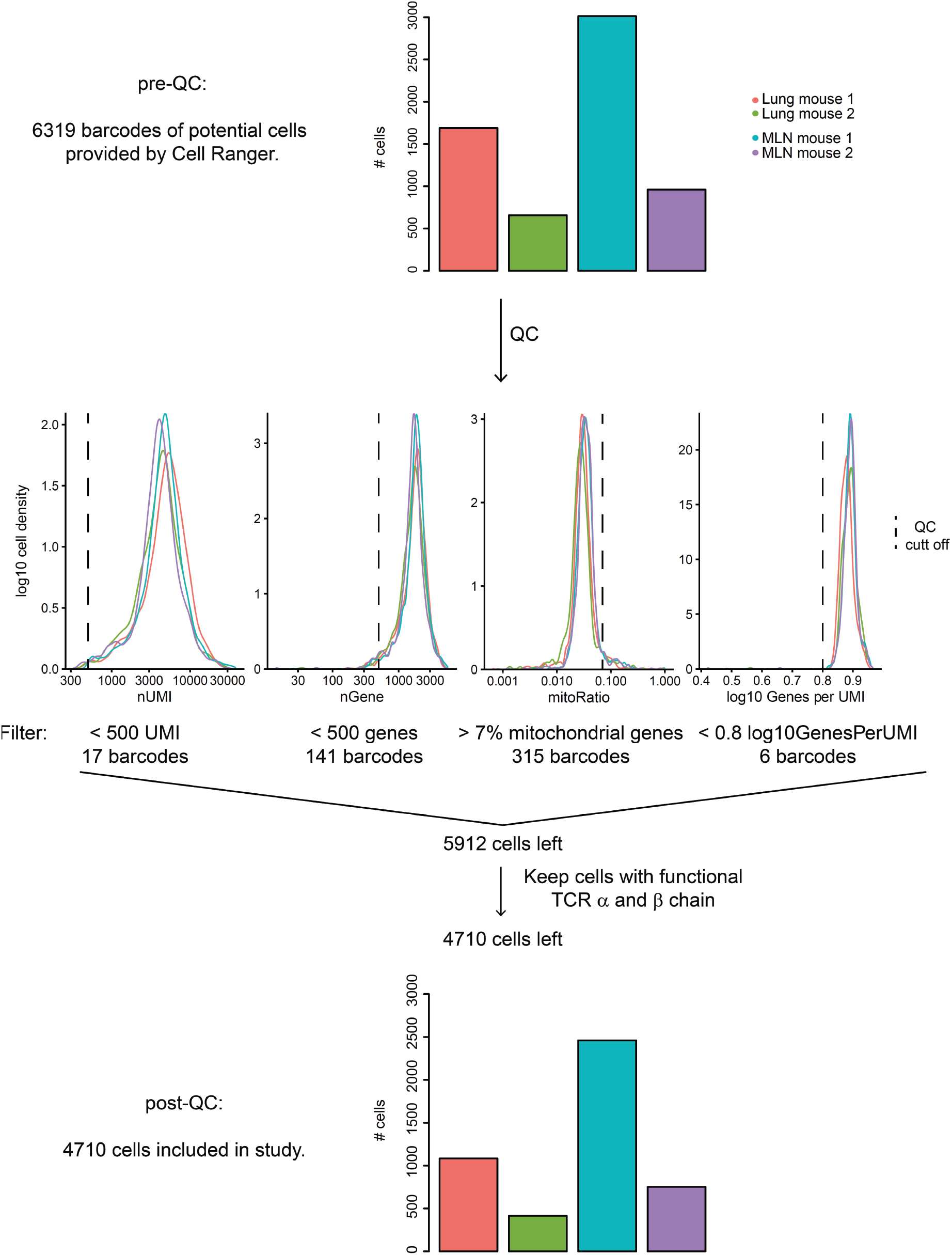
Quality control of Th2 single cell sequencing ten days post Nb infection. Overview of QC workflow. Numbers of potential cells pre-QC are given for different mice and organs (upper panel). Histograms visualize exclusion of potential cells by various cut offs (middle). Numbers of included cells post QC are visualized for different mice and organs (lower panel). Functional TCR chains according to Cell Ranger definition.

**Suppl. Figure 2.**
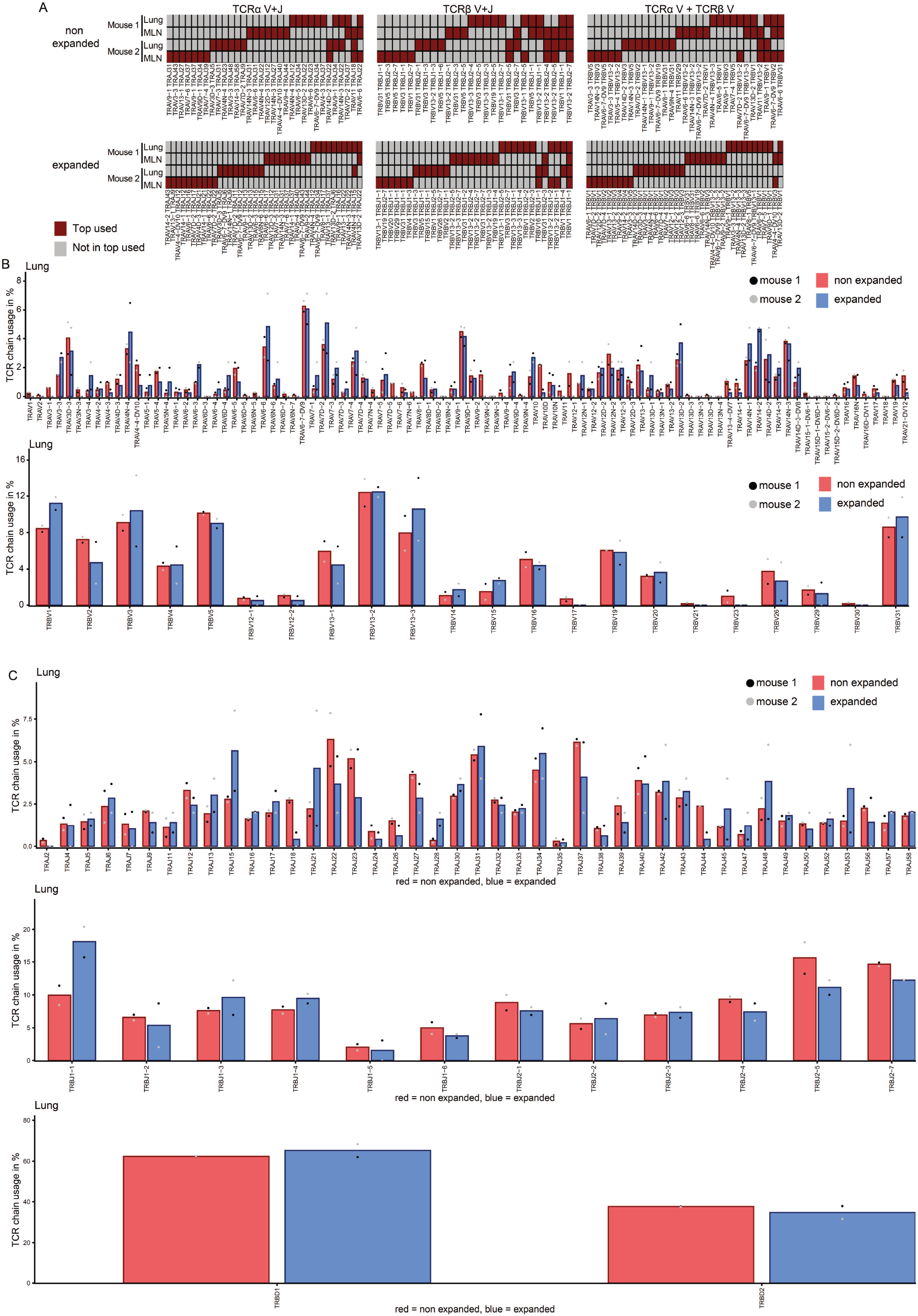
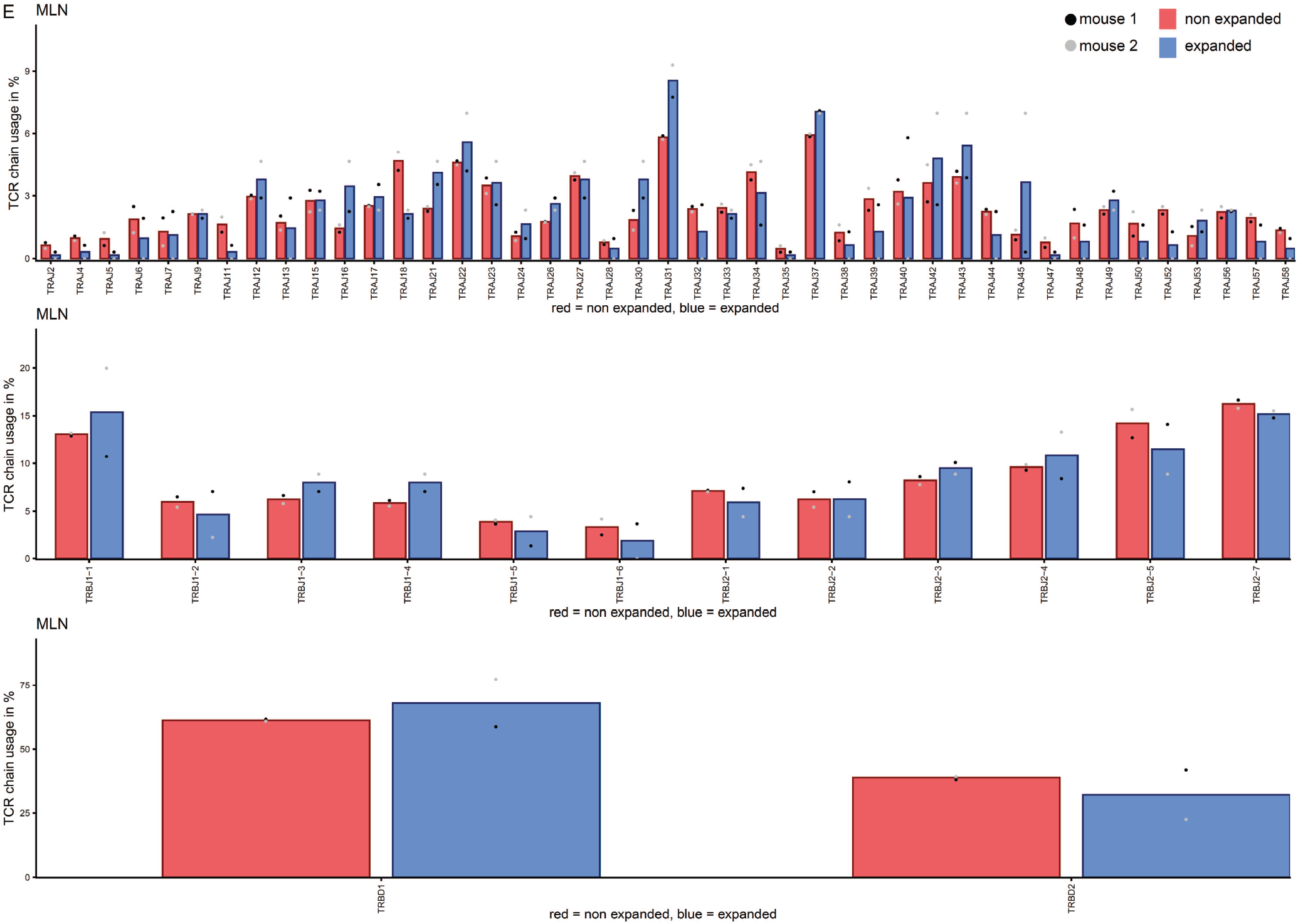
TCR repertoire analysis of MLN and lung Th2 cells at day ten post Nb infection. A) Overlap of functional TCRs that use the same combination of TCRα V+J segments (left), TCRβ V+J segments (center), or TCRα V + TCRβ V segments (right) among the ten top used combinations for each sample. This was determined separately for expanded and non-expanded clones. One cell of each clone was considered to avoid expansion bias. Changes in variable regions were not further taken into account for overlap determination. B) Usage of different TCRα variable or TCRβ variable segments in lung Th2 cells. C) Usage of different TCRα joining, TCRβ joining and diversity segments in lung Th2 cells. D) and E) as B) and C) but for MLN.

**Suppl. Figure 3.**
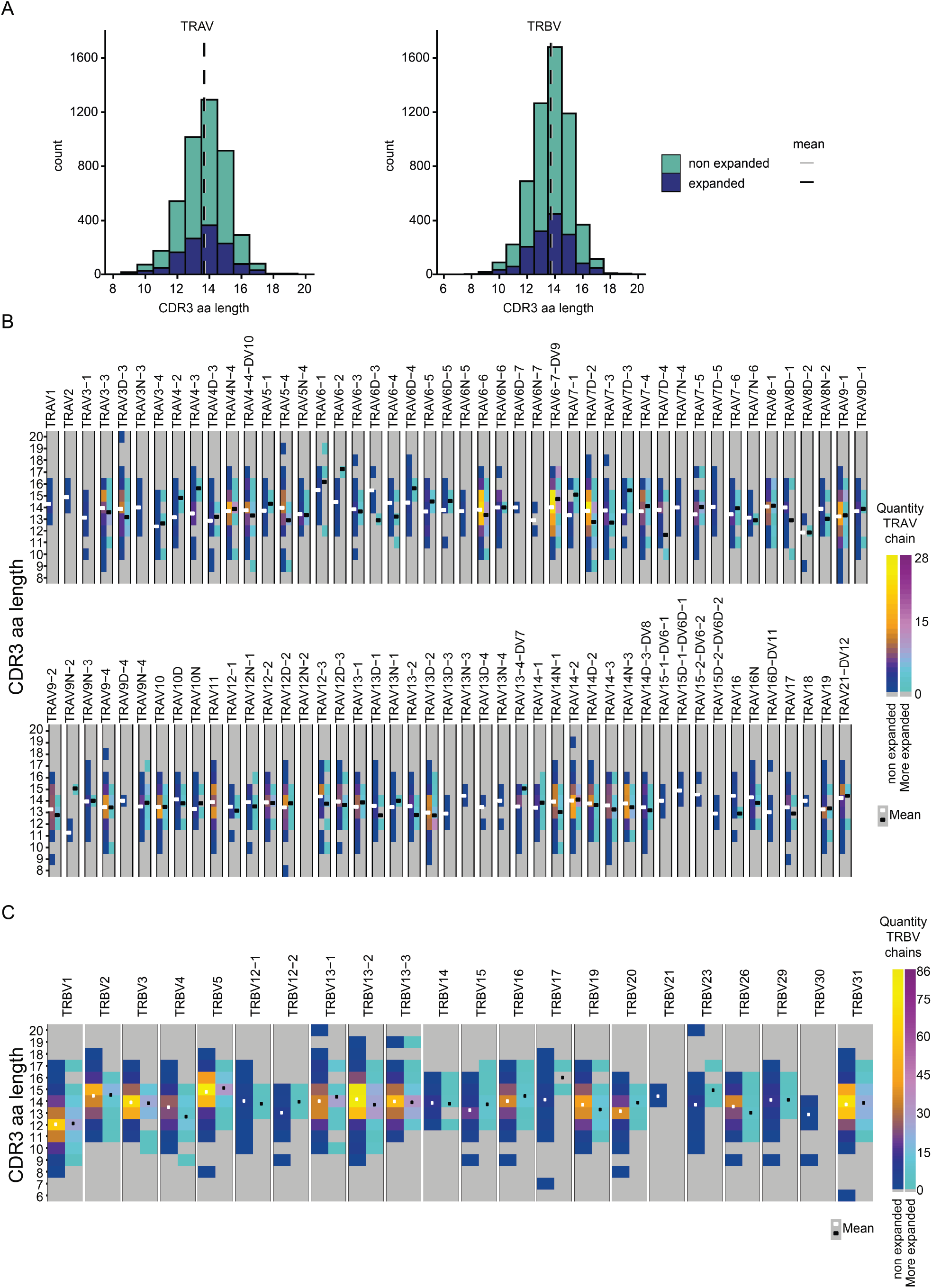
CDR3 length of Th2 TCRα and TCRβ chains ten days post Nb infection. A) TCRα and TCRβ CDR3 amino acid length. Data of MLN and lung cells were combined. CDR3 length was determined separately for expanded and non-expanded clones. One cell of each clone was considered to avoid expansion bias. B) CDR3 amino acid length distribution for all used TCRα variable segments or C) all used TCRβ variable segments.

